# Sex-specific role of the circadian transcription factor NPAS2 in opioid tolerance, withdrawal, and analgesia

**DOI:** 10.1101/2022.03.27.485966

**Authors:** Stephanie Puig, Micah A. Shelton, Kelly Barko, Marianne L. Seney, Ryan W. Logan

## Abstract

Opioids like fentanyl are the mainstay treatment for chronic pain. Unfortunately, opioids high addiction liability has led to the current opioid crisis. This is in part related to long-term use side-effects, including analgesic tolerance (gradual decrease in analgesia), and physical dependence (withdrawal symptoms upon opioid interruption), causing dose-escalation and preventing usage interruption. Altered circadian rhythmicity and sleep patterns are common in patients on opioid therapy. Prior research showed intricate bidirectional interactions between circadian rhythmicity, opioid analgesic efficacy, and side-effects. However, underlying mechanisms are largely unknown. Neuronal PAS domain protein 2 (NPAS2) is a circadian transcription factor that is highly expressed in structures of the central nervous system that modulate pain and opioids. In this study, we used male and female mice expressing non-functional NPAS2 (NPAS2 deficient, NPAS2-/-) to investigate the role of NPAS2 in fentanyl analgesia, tolerance, hyperalgesia, and dependence. In NPAS2-/- mice, we found that thermal pain thresholds, acute analgesia, and tolerance to a fixed dose of fentanyl were largely like wild-type mice. However, female NPAS2-/- mice augmented behavioral state of analgesic tolerance and exhibited significantly more behavioral symptoms of physical dependence. Conversely, only male NPAS2-/- mice had increased fentanyl-induced hypersensitivity, when compared to matched-sex littermate controls. Together, our findings suggest sex-specific effects of NPAS2 signaling in the regulation of fentanyl-induced tolerance, hyperalgesia, and dependence.

## INTRODUCTION

Prescription opioids are potent analgesics widely used to treat pain. Extended use, often necessary to treat chronic pain, is a risk factor for developing tolerance (*e.g*., reduced effect of analgesia), physical dependence (*i.e*., withdrawal symptoms during periods of abstinence), and pain hypersensitivity (*i.e*., opioid-induced increases in pain sensitivity) ^1,2^, and serves as a risk factor for opioid use disorder ^3^. Hallmarks of opioid dependence and opioid use disorder are profound disruptions to sleep and circadian rhythms that persist during abstinence and withdrawal ^4-7^. In line with this, previous evidence suggests opioid tolerance, dependence, and analgesia ^8-12^ are modulated by sleep and circadian rhythms. Altered circadian rhythms and sleep disruptions are commonly observed in patients being treated with opioids ^13^. For example, sleep disturbances are a known indicator of opioid withdrawal symptoms in patients attempting to discontinue their opioid treatment ^14^. Sleep and circadian rhythm disruptions may also contribute to relapse risk in opioid abstinent patients ^15,16^. However, our understanding of the molecular pathways that underlie the relationship between circadian rhythms and opioid tolerance and withdrawal is limited.

Nearly every cell in the body expresses the necessary machinery that controls circadian rhythms at the molecular level^17^. The molecular clock is composed of a series of interacting transcriptional and translational feedback loops. Transcription is driven by heterodimers of CLOCK (Circadian Locomotor Output Cycles Kaput Protein), or NPAS2 (Neuronal PAS domain protein 2), with BMAL1 (brain muscle aryl nuclear translocase like-1), that bind to enhancer elements in gene promoters. These heterodimers of circadian transcription factors (CLOCK or NPAS2 and BMAL1) drive the transcription of circadian genes, *Cry1,2* (*Cryptochrome*) and *Per1,2* (*Period*). Accumulation of CRYs and PERs in the cytoplasm over 24-hours initiate the formation of their own dimers that eventually translocate to the nucleus to inhibit their own transcription via interactions with BMAL1 heterodimers ^18^. Molecular clocks are critical to the function of most peripheral organs and many regions in the brain ^19^.

In peripheral tissues and the central nervous system, the molecular clock modulates the circadian expression of endogenous opioid peptides and opioid receptors ^20-23^. Opioids also impact the expression of circadian genes in the brain. For example, acute and chronic administration of opioids alters circadian genes in the hypothalamus ^24-26^, while withdrawal from opioids alters the rhythmic expression of circadian genes in the midbrain and striatum ^26-28^. Recent findings demonstrate circadian genes, including *Per1* and *Per2*, regulate opioid reward ^29^, tolerance, and withdrawal ^30^. In a tissue-dependent manner, *Per1* and *Per2* transcription are driven by BMAL1 dimerization with CLOCK or NPAS2 ^31-33^. Notably, the circadian transcription factor NPAS2 is enriched in major neural substrates of opioid-induced tolerance ^34^, dependence ^35-37^, and hyperalgesia ^38^, including the spinal cord ^39^, primary sensory neurons ^40^, and within the subregions of the striatum, including the nucleus accumbens (NAc) ^41^. However, to our knowledge, the involvement of NPAS2 signaling in the emergence of opioid side-effects has not yet been studied.

In the present study, we investigated the role of NPAS2 in opioid tolerance, dependence, and analgesia, using the synthetic opioid, fentanyl. Fentanyl is a widely prescribed opioid with high addiction potential, found in most drug overdose deaths in the United States ^3^. Thus, further investigation into the potential pathways mediating the development of fentanyl-induced tolerance and dependence is imperative, as new therapeutic approaches are designed to improve opioid treatments and mitigate secondary effects of opioids. To explore the potential relationship between NPAS2 and opioids, we assayed behavioral phenotypes of fentanyl-induced tolerance, dependence, and hyperalgesia in NPAS2 deficient male and female mice (NPAS2-/-). These transgenic mice possess a LacZ reporter in replacement of the basic Helix-loop-Helix (bHLH) domain required for NPAS2 to directly bind DNA and regulate transcription ^32^. Therefore, these NPAS2-/- mice retain the expression of NPAS2 protein, while lacking the ability to bind DNA and drive transcription. Overall, our findings reveal sex-specific effects of NPAS2 deficiency on thermal nociceptive thresholds, acute anti-nociception, and development of tolerance to fentanyl, as well as behavioral markers of physical dependence and pain hypersensitivity.

## METERIALS & METHODS

### Animals

Adult male and female NPAS2 deficient (NPAS2-/-) mice and their respective wild-type (WT) littermates were used (aged 7-14 weeks) ^32^. Initial breeders were generously provided by Dr. David Weaver (University of Massachusetts Medical School). NPAS2-/- mice have a LacZ-Neo fusion gene in inserted into exon 2, replacing the locus that encodes the bHLH domain. Replacing the bHLH domain renders the NPAS2 protein deficient of capability to directly bind DNA and mediate NPAS2-dependent gene transcription. Mice were maintained on C57BL6/J background (The Jackson Laboratory, 000664; backcrossed to at least N10). Mice were group housed (2-4 mice per cage) and maintained under 12h:12h standard light-dark cycle with *ad libitum* access to food and water. Experimental procedures were approved by the Institutional Animal Care and Use Committee at the University of Pittsburgh School of Medicine.

### Drugs

Fentanyl hydrochloride (NIH NIDA, Bethesda, MD) and Naloxone (Sigma, St Louis, MO) were dissolved in sterile saline vehicle (0.9%) under a biosafety cabinet. Fentanyl was administered intraperitoneally (i.p., 10ml/kg, v/w).

### General Procedures and Experimental Timeline

Throughout the experiments, experimenters were blind to both the treatment and genotype groups. Prior to the each of the behavioral assays, mice were habituated to experimental rooms for at least one hour per day for three consecutive days. During habituation, mice were gently handled by the experimenters to prepare for various procedures required to conduct the behavioral assays. For the behavioral assays 9 to 11 mice per genotype per used. The experimental timeline is described in **Figure 1**. On day 0, baseline thermal thresholds were measured from tail flick latencies (TFL) averaged over three trials per mouse. On day 1, mice underwent a first dose-response procedure with fentanyl (i.p.). TFLs were measured 15 mins after fentanyl administration. The fentanyl dose was then doubled every 20 mins, and tested again until TFLs reached the threshold of 10 sec. The following fentanyl doses were administered on day 1: 10, 20, 40, 80, 160 and 320 μg/kg. Between days 4 through 8, mice were tested for the development of tolerance and hyperalgesia. Mice were administrated fentanyl (320 μg/kg, i.p.) twice daily at ∼ZT2 (900h, lights on at 700h) and ∼ZT9 (1600h). Testing of TFL was performed before hyperalgesia and 15 minutes after tolerance testing at the ZT2 administration. On day 11, mice a second dose-response was conducted using the following fentanyl doses: 160, 320, 640 and 1280 μg/kg. Finally, on day 12, mice underwent naloxone-precipitated withdrawal. Mice were first administered fentanyl (320 μg/kg, i.p.), then ∼15 mins later, naloxone (10mg/kg, i.p.). Mice were immediately placed in a novel cage and withdrawal behaviors were recorded.

**Figure 1.**
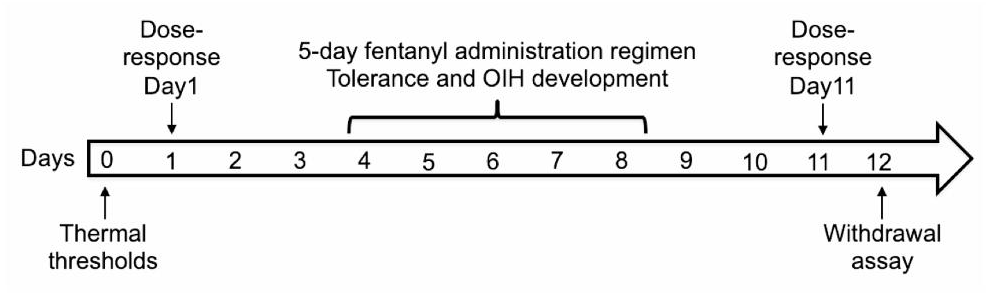
Experimental timeline. All animals underwent the same procedures. On day 0, animals were tested for thermal nociceptive thresholds (baselines) with the tail flick (TFL) assay. On day 1, a first dose-response of the fentanyl analgesic effect on TFL was performed. The 5-day fentanyl administration regimen involving two fentanyl i.p. injections per day was conducted from day 4 to day 8. Tolerance and OIH development were evaluated simultaneously. On day 11, a second fentanyl dose-response was conducted. This series of experiments concluded with the naloxone-precipitated withdrawal assay on day 12.

### Tail-immersion test for fentanyl-induced analgesia and tolerance

The tail-immersion behavioral assay was used to assess thresholds of thermal nociception, analgesia, and tolerance to fentanyl. A container filled with water maintained at 50±0.5°C using a thermoregulated water bath. Part of the mouse’s tail was immersed in the water and the TFL was recorded with a maximum immersion limit of 10 seconds to avoid tissue damage. After tasting, mice were immediately returned to their home cage. TFL measurements were repeated 2-3 times per mouse at 2 mins intervals.

### Naloxone-precipitated fentanyl withdrawal

Following the development of physical dependence to fentanyl, behavioral signs of withdrawal from chronic fentanyl were evaluated using the opioid receptor antagonist, naloxone. Mice were administered fentanyl (320 μg/kg, i.p.), then naloxone (10mg/kg, i.p.) ∼15 mins later. Withdrawal behaviors were recorded for 10 mins in a novel home cage, which included escape jumps, wet-dog shakes, paw-shakes/grooming, teeth chattering. Presence of diarrhea was also recorded. Behaviors were compiled to calculate cumulative withdrawal scores as follows: Jumps: 1-10 = 2, 11-20 = 4, 21-30 = 6, 31-40 = 8, 41-50 = 10, etc.; wet dog shakes: 1-2 = 2, 3-10 = 4, 10 and more = 6; and presence of teeth chattering, paw shakes and diarrhea were scored a 2.

### Statistical analyses

A combination of statistical analyses was used for the behavioral assays. Tolerance measures were analyzed using two-way mixed effect ANOVA (Time and Treatment) followed by Tukey’s multiple comparison *post-hoc* analyses. Thermal nociceptive thresholds, EC50s and withdrawal behaviors were analyzed using one-way ANOVA, followed by Tukey’s post-hoc multiple comparison tests, when appropriate. Dose-response curves were generated using a non-linear regression of log-transformed values compared to normalized values from 0 to 100. Comparisons of rightward shifts between mouse genotypes were done by calculating the ratio between EC50 obtained on day 11 / EC50 obtained on day 1. Unpaired t-test comparisons were performed to determine difference of rightward shift between NPAS2-/- and WT mice of the same sex. Data were analyzed using GraphPad 9.0 and considered statistically significant if p≤0.05.

## RESULTS

### Impact of NPAS2 deficiency on thermal thresholds, fentanyl acute analgesia, fentanyl-mediated tolerance, and hypersensitivity

The impact of NPAS2 deficiency on thermal thresholds and development of tolerance and hypersensitivity with chronic fentanyl, was evaluated by measuring Tail Flick Latencies (TFLs) in NPAS2-/- and WT mice before and after a 5-day fentanyl administration regimen (**Figure 1**). The acute analgesic effect of fentanyl measured on day 1 was similar between NPAS2-/- and WT male (**Figure 2A**) and female (**Figure 2B**) mice. Additionally, all groups had a similar progressive reduction in TFLs over time without a significant difference between genotypes in males (**Figure 2A**, 2-way ANOVA, Interaction: F_5, 90_ = 0.1601, P = 0.9764, Days: F_3.058, 55.04_ = 48.38, P < 0.0001, Treatment: F_1, 18_ = 0.7066, P < 0.0001) or females (**Figure 2B**, 2-way ANOVA, Interaction: F_5, 80_ = 2.233, P = 0.0590, Days: F_3.211, 51.37_ = 50.40, P < 0.0001, Treatment: F_1, 16_ = 0.001806, P < 0.9666). Similarly, baseline TFLs measured in opioid naïve animals prior to fentanyl administration, were largely similar between NPAS2-/- and WT male and female mice **(Figure 2C**), suggesting that NPAS2 has minimal to no measurable impact on thermal nociceptive thresholds in opioid naïve mice (1-way ANOVA, F_3, 34_ = 0.8418, P = 0.4805). However, thermal thresholds measured prior to fentanyl administration on day 1 compared to day 5 revealed NPAS2-/- male mice developed thermal hypersensitivity, an effect not seen in WT male mice. Conversely, both NPAS2-/- and WT females developed thermal hypersensitivity (**Figure 2D**, day 5 – day 1 TFLs, F_3, 34_ = 2.765, P = 0.0568). Together, these findings support the possible involvement of NPAS2 in hyperalgesia development in a sex-specific manner.

**Figure 2.**
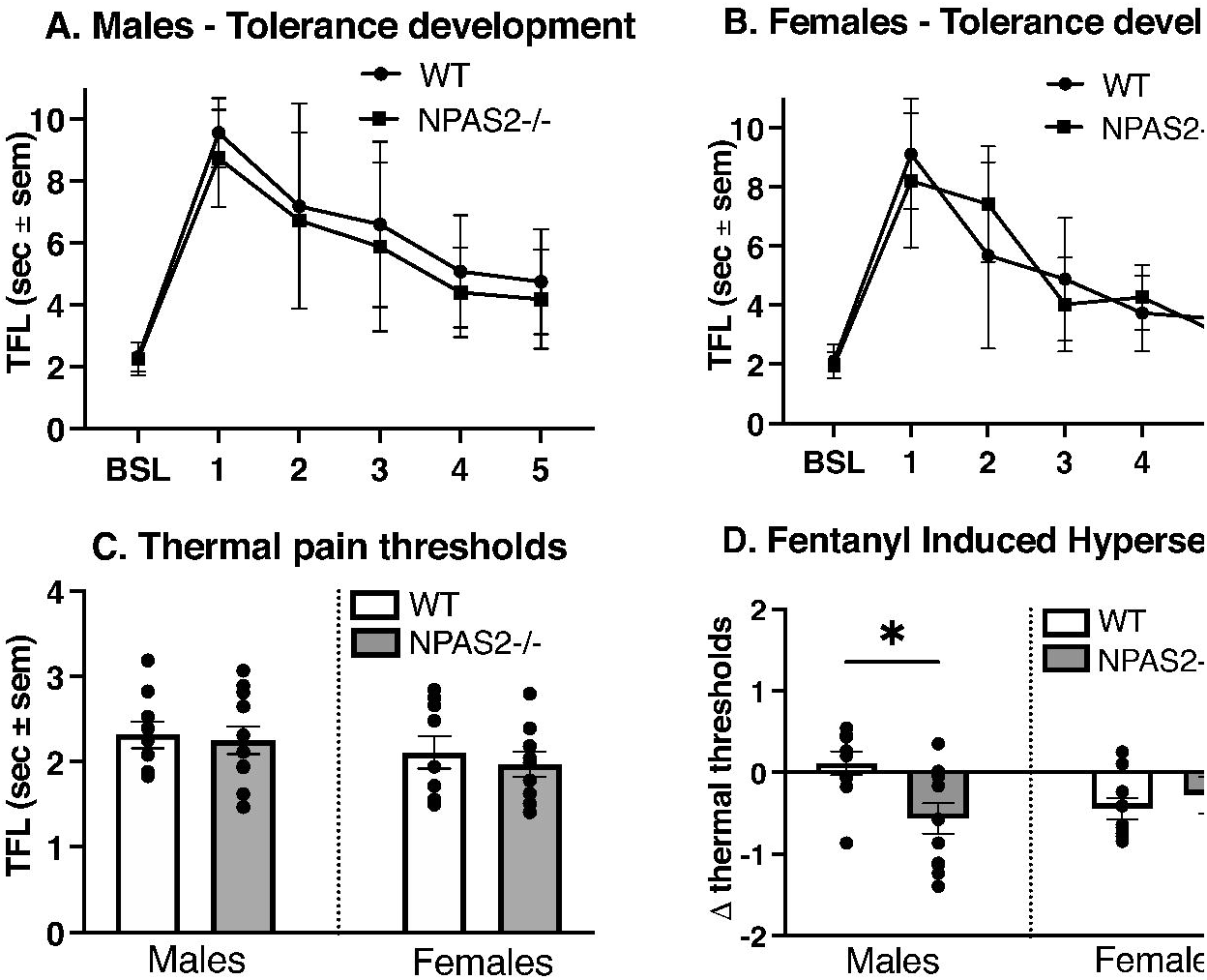
NPAS deficiency does not alter thermal nociceptive thresholds or fentanyl analgesic tolerance development but promotes fentanyl-induced hypersensitivity in males only. Development of tolerance to a fixed dose of fentanyl (320μg/kg) administered twice a day for 5 days was assessed my measuring tail flick latencies (TFLs), daily in males **(A)**, and females **(B)**. Two-way ANOVAs, males, N = 9-11, Interaction: F_5, 90_ = 0.1601, *p* = 0.9764, Days: F_3.058, 55.04_ = 48.38, *p* < 0.0001, Treatment: F_1, 18_ = 0.7066, *p* < 0.0001; females, N = 11, Interaction: F_5, 80_ = 2.233, *p* = 0.0590, Days: F_3.211, 51.37_ = 50.40, *p* < 0.0001, Treatment: F_1, 16_ = 0.001806, *p* < 0.9666. **(C)** Baseline thermal nociceptive thresholds measured prior to the beginning of fentanyl injections. One-way ANOVA, N = 9-11, F_3, 34_ = 0.8418, *p* = 0.4805. **(D)** Comparison of baseline thermal threshold measured on day 0, to threshold measured on day 5 before the fentanyl injection. Data represented as delta of TFL values measured on Day 5 – Day 0, 1-way ANOVA, N = 9-11, F_3, 34_ = 2.765, *p* = 0.0568. Data represented as mean +/-SEM. TFL: tail flick latency, BSL: Baseline.

### Impact of NPAS2 deficiency on changes in fentanyl potency due to tolerance

To further test the impact of NPAS2 deficiency on fentanyl analgesic potency, we performed two dose-response procedures on male and female NPAS2-/- and WT mice. The first dose-response was conducted on day 1 when mice were naïve to opioids (**Figure 3A)**, and the second dose-response was conducted on day 11, after mice had received the tolerance-inducing 5-day fentanyl administration regimen (**Figure 3B**). Dose-responses performed on day 11, revealed a rightward shift of the fentanyl dose-response curves in all groups, as compared to dose-responses measured on day 1 (Males, **Figure 3A**, Females, **Figure 3B**). Therefore, our regimen induced a robust state of analgesic tolerance in all mice. Based on best-fit values of these curves, we calculated effective concentration 50 (EC50) values, which represent the concentration of fentanyl that gives half-maximal analgesia, (EC50 day 1 values: WT males: 51.75 μg/kg; NPAS2-/- males: 61.25 μg/kg; WT females: 71.24 μg/kg; and NPAS2-/- females: 63.01 μg/kg. EC50 day 11 values: WT males: 273.4 μg/kg, NPAS2-/- males: 269.6 μg/kg, WT females: 291.0 μg/kg and NPAS2 females: 346.0 μg/kg). For EC50 values, no significant differences between NPAS2-/- and WT mice were found in pre-tolerance fentanyl potency on day 1 (**Figure 3C**, one-way ANOVA, F_3,33_ = 0.4501, P = 0.7189). On day 10, EC50 values were also similar between genotypes and among both sexes (**Figure 3D**, 1-way ANOVA, F_3,33_ = 2.056, P = 0.1251). Finally, calculation of the rightward shift factor between naïve (day 1) and tolerant (day 11) mice (EC50 day1 – EC50 day11), revealed that degree of rightward shift of curves in WT and NPAS2-/- males were similar (**Figure 3E**, two-tailed t-test, t=0.6714, df=18, p=0.5105). Conversely, comparison of shift factors in female mice revealed a greater shift in NPAS2-/- females (6.069 rightward shift average), than in WT females (4.382 rightward shift average) that was close to statistical significance (**Figure 3F**, two-tailed t-test, t=2.096, df=16, p=0.0523). Overall, our findings point toward a possible impact of NPAS2 deficiency on fentanyl potency in female, but not male, mice.

**Figure 3.**
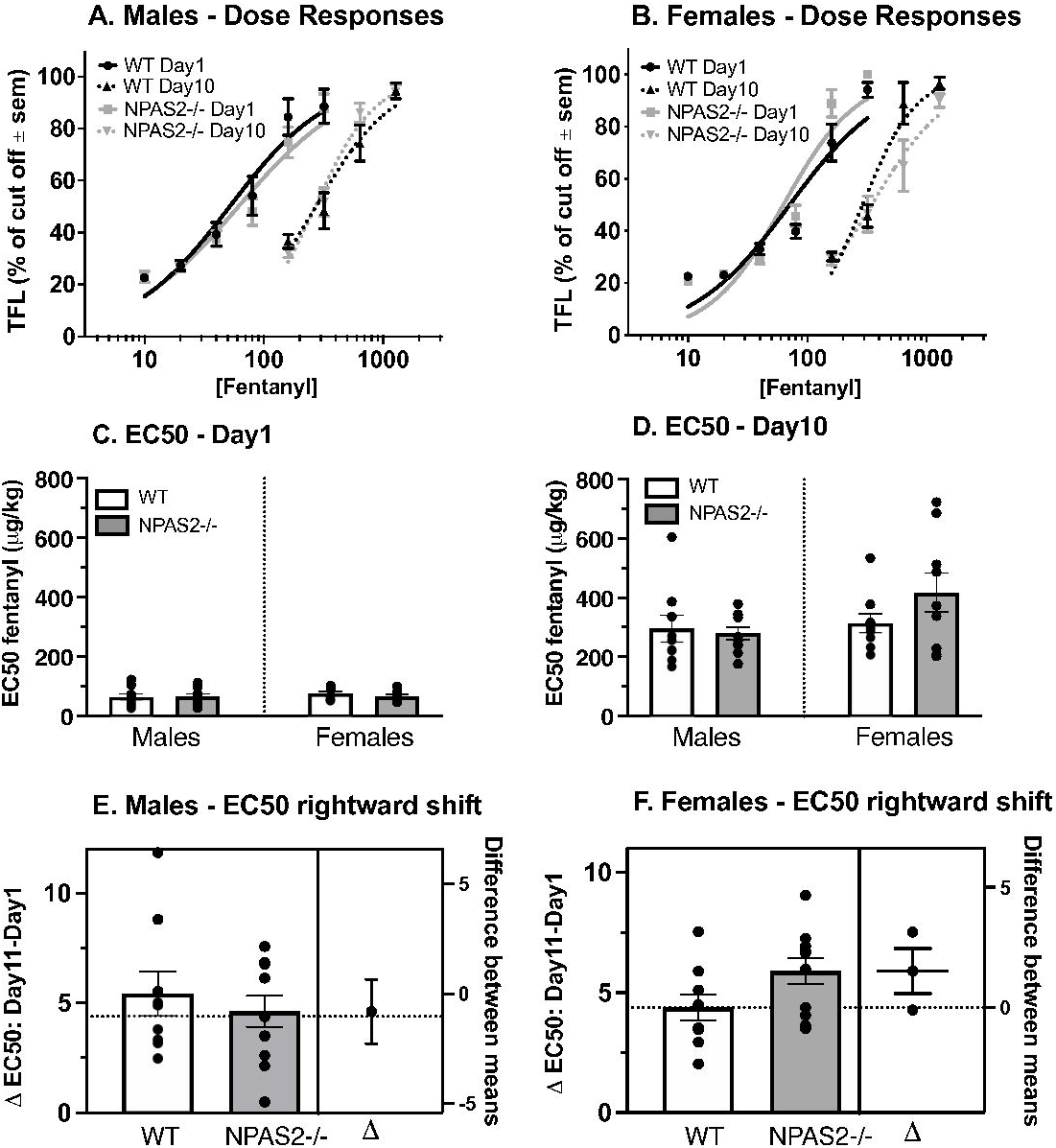
NPAS2-deficiency alters fentanyl potency in female tolerant mice. Fentanyl dose-response curves in males **(A)**, and females **(B)** measured before (day1) and after (day11) induction of tolerance with fentanyl. Doses administered ranged from 10μg/kg to 320μg/kg for during the pre-tolerance test, and from 160μg/kg to 1280μg/kg during the post-tolerance test. Data represented as normalized log[fentanyl] and normalized from 0 to 100. N = 9-11, Non-linear fit, Best-fit values for EC50 values calculations: Males: WT pre-tolerance: 51.75 μg/kg, WT post-tolerance: 273.4 μg/kg, 5.1-fold rightward shift; NPAS-/- pre-tolerance: 61.25 μg/kg, NPAS-/- post-tolerance: 269.6 μg/kg, 4.4-fold rightward shift. Females: WT pre-tolerance: 71.24 μg/kg, WT post-tolerance: 291.0 μg/kg, 4.08-fold rightward shift; NPAS-/- pre-tolerance: 63.01 μg/kg, NPAS-/- post-tolerance: 346.0 μg/kg, 5.5-fold rightward shift. **(C)** Day1 EC50s comparisons, N = 9-10, 1-way ANOVA, F_3,33_ = 0.4501, *p* = 0.7189. **(D)** Day11 EC50s comparisons, N = 9-10, 1-way ANOVA, F_3,33_ = 2.056, *p* = 0.1251. **(E)** Comparison of degree of rightward shift of EC50s between day1 and day11 (EC50 day1-EC50 day11) degree between WT and NPAS2-/- males, N = 9-11, two-tailed t-test, t = 0.6714, df = 18, *p* = 0.5105. **(F)** Comparison of degree of rightward shift of EC50s between day1 and day11 (EC50 day1-EC50 day11) degree between WT and NPAS2-/- females, N = 9, two-tailed t-test, t = 2.096, df = 16, *p* = 0.0523. **(C-D)** Data represented as mean +/- SEM. TFL = tail flick latency, EC50 = effective dose 50.

### Impact of NPAS2 deficiency on naloxone-precipitated withdrawal responses in fentanyl-dependent mice

Following the post-tolerance dose-response regimen of fentanyl, mice received a fentanyl challenge, followed by naloxone, to precipitate withdrawal and induce dependence behaviors. Overall, female NPAS2-/- mice displayed significantly more withdrawal behaviors than NPAS2-/- and WT male mice. Fentanyl naloxone precipitated withdrawal led to more jumps in NPAS2-/- females compared to WT females (**Figure 4A**, 1-way ANOVA, F_3, 34_ = 4.646, P = 0.0079), while these behaviors were similar between genotypes in males. The number of wet-dog shakes were overall unchanged in NPAS2-/- mice (**Figure 4B**), although female mice had significantly more wet-dog shakes than males, regardless of genotype (**Figure 4B**, 1-way ANOVA, F_3, 34_ = 3.425, P = 0.0279). In addition, teeth-chattering episodes were more frequent in NPAS-/- mice as compared to WT mice, but this effect was not significant (**Figure 4C**, 1-way ANOVA, F3, 34 = 2.624, P = 0.0663). Finally, no difference was observed in the number of grooming/paw shaking episodes across groups (**Figure 4D**, 1-way ANOVA, F_3, 34_ = 0.8478, P = 0.4774). Global withdrawal scores, which involved the number of jumps, along with the number of teeth-chattering, wet dog shakes, grooming/paw shakes episodes, showed that NPAS2-/- females displayed significantly more withdrawal behaviors as compared WT female mice and male mice from both genotypes (**Figure 4E**, 1-way ANOVA, F_3, 34_ = 6.543, P = 0.0013.). Together, these results support a role of NPAS2 in the development and exhibition of physical dependence primarily in female than in male mice.

**Figure 4.**
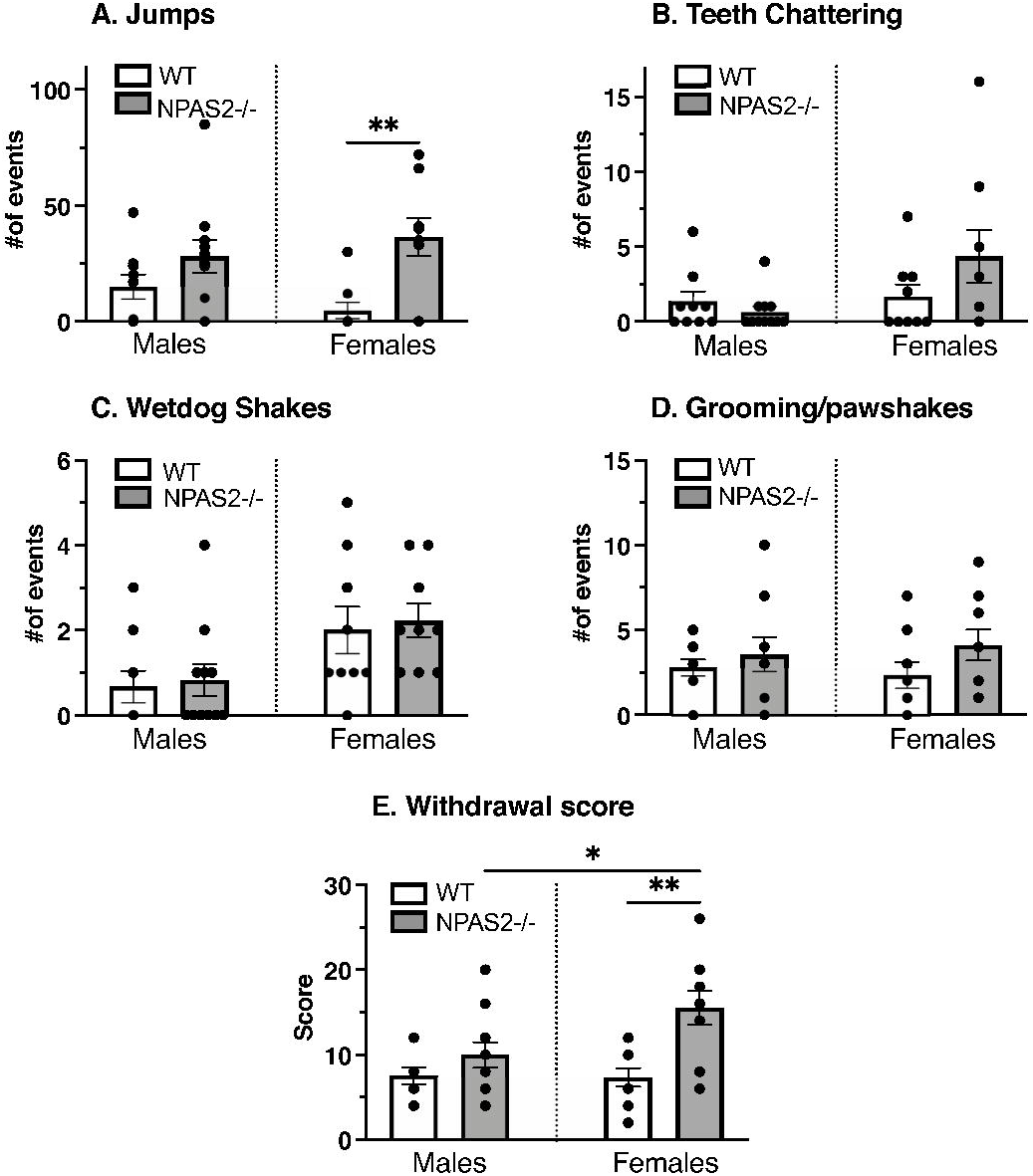
NPAS2-deficiency potentiates physical dependence behaviors in female mice only. Naloxone-precipitated withdrawal behaviors in NPAS2-/- and WT littermate mice administered with a challenge dose of fentanyl (320μg/kg, i.p.). **(A)** total number of jumps, N = 9-11, 1-way ANOVA, F_3, 34_ = 4.646, *p* = 0.0079. **(B)** total number of wet dog shake episodes, N = 9-11, 1-way ANOVA, F_3, 34_ = 3.425, *p* = 0.0279. **(C)** total number of teeth chattering episodes, N = 9-11, 1-way ANOVA, F_3, 34_ = 2.624, *p* = 0.0663. **(D)** total number of paw shakes/grooming episodes, N = 9-11, 1-way ANOVA, F_3, 34_ = 0.8478, P = 0.4774. **(E)** withdrawal score, N = 9-11, 1-way ANOVA, F_3, 34_ = 6.543, *p* = 0.0013. Data represented as mean +/- SEM. Tukey’s multiple comparisons tests, * *p* < 0.05, ** *p* < 0.01, *** *p* < 0.001.

## DISCUSSION

Long-term opioid use for the treatment of chronic pain, is hampered by analgesic tolerance^42^, physical dependence^43^, and profound disruptions of sleep and circadian rhythms ^4-7^. Previous studies implicated circadian regulation of pain, tolerance, and physical dependence ^11,12^, suggesting bidirectional interactions between circadian rhythms and opioids ^9^. NPAS2 is a circadian gene enriched in brain regions and spinal cord involved in pain and opioids, and previously shown to be involved in psychostimulant reward ^44,45^. However, the role of NPAS2 in opioid tolerance and dependence had yet to be established. To address this, we used NPAS2-deficient mice to investigate the involvement of NPAS2 on fentanyl-mediated tolerance, hypersensitivity, and physical dependence. Overall, we found that thermal pain thresholds, acute fentanyl analgesia, and fentanyl tolerance development were unchanged in male and female NPAS2-/- mice. Interestingly, NPAS2 deficiency led to a decrease in fentanyl potency after tolerance developed and led to markedly more symptoms of physical dependence only in female mice. Conversely, NPAS2 deficiency was associated with increased fentanyl-induced hypersensitivity in male compared to female mice.

Our results are consistent with prior studies which also evaluated the impact of genetic deletion of circadian genes on pain and opioids ^29,30^. In these studies, mice with global deletion of *mPer1* (mPer1-KO)^29^ or *mPer2* (mPer2-KO)^30^ genes showed no change in thermal pain thresholds and acute analgesia. This is similar to our observations with NPAS2-/- mice and suggests that these circadian genes may not be directly involved in these behaviors. Conversely, *mPer1* and *mPer2* clock genes were differentially involved in tolerance and dependence, as mPer2-KO promoted morphine tolerance and mitigated physical dependence^30^, while mPer1*-*KO did not affect tolerance or physical dependence^29^. In our study, which included both males and females, NPAS2-/- mice did show altered tolerance development to a fixed dose of fentanyl across both sexes. In addition, we also evaluated fentanyl potency, by conducting dose-response tests before and after tolerance development. Strikingly, NPAS2-/- mice developed higher analgesic tolerance compared to controls, an effect only observed in females. Together, these data could suggest that circadian genes are differentially involved in tolerance, with *mPer2* being involved in its development, *Npas2* in its expression, and *mPer1* likely not involved. However, caveats of studies investigating the roles of *Per* genes include the absence of testing opioid potency in tolerant animals with a dose-response assay, along with these studies including only male mice. Because *Npas2* and *Per* genes are both expressed in structures involved in tolerance such as the spinal cord ^39,46^ or the NAc^47^, and NPAS2 can directly regulate the transcription of *Per* genes, involvement of each of these circadian genes in tolerance remains a possibility.

Sex specific effects of NPAS2 deficiency were also observed with physical dependence symptoms. NPAS2-/- females showed more withdrawal behaviors than controls, also more than NPAS2-/- males. Conversely, when testing fentanyl-induced hypersensitivity, a symptom also emerging during opioid withdrawal, NPAS2-/- females were similar to WT females, while NPAS2-/- males had more pronounced hypersensitivity than WT males. Together, these results suggest that tolerance, physical dependence, and opioid-induced hypersensitivity behaviors could be modulated by NPAS2 signaling in a sex specific manner. Involvement of NPAS2 in these behaviors could be supported by the fact that NPAS2 expression is enriched in the NAc ^32,41^. The NAc is involved in tolerance^47,48^, physical dependence^36,49,50^, and opioid-induced hyperalgesia^12^. NPAS2 modulates dopaminergic and glutamatergic neurotransmission in the striatum ^51^, both of which are altered during opioid tolerance ^47,48^ and withdrawal ^52-54^. Interestingly, changes in the expression of circadian genes in the NAc were shown to occur in rodents with opioid-induced hyperalgesia during a state of withdrawal ^12^. Thus, together with our current findings, NPAS2 may modulate opioid-related behaviors and involve dopaminergic and glutamatergic signaling in the NAc. Ongoing and future studies are exploring possible involvement of NPAS2 in the NAc and the dopaminergic and glutamatergic neurotransmission in opioid tolerance and dependence.

Importantly, our results illustrate the importance of performing these further including both males and females to examine intersectional consequences between sex and opioid effects, and between sex and circadian genes. This is also supported by several previous studies, which examined whether sex could have a differential impact on opioid behaviors. Analgesic effect of opioids was shown to be variable depending on sex, with a higher and longer-lasting effect in male than female rodents ^55-57^, although, other studies did not find a sex difference in opioid analgesia ^58,59^. In addition, sexual dimorphism in opioid tolerance has not been extensively studied in rodents, yet the studies that have, report higher tolerance in males than females ^60^, with tolerance developing faster in females than males ^61,62^. However, these findings have not been supported by other studies ^63-65^. Finally, sexual dimorphism in opioid dependence-mediated withdrawal behaviors were also reported in rodent studies, with overall more dependence in WT males than WT female rats ^60,66^. Inconsistent findings between men and women have also been reported in humans ^67-70^. In our current study, we did not observe sex-related differences in opioid analgesia, tolerance development and expression, dependence, and hyperalgesia, between WT male and female mice. Together with these aforementioned studies, our data illustrates the lack of consensus on the impact of sex on opioid behaviors, and thus require further investigation.

Nevertheless, in our study, we specifically examined intersectional consequences of NPAS2 deficiency and sex on opioid analgesia, tolerance, and dependence behaviors. Interestingly, NPAS2-/- females developed markedly more physical dependence behaviors and marginally more profound tolerance than female WT littermates, which was not observed in males. However, NPAS2 deficiency had no consequences on hyperalgesia development in females while it exacerbated that symptom in males. Thus, our data indicates that sex differences in our study are related to interacting effects between NPAS2 deficiency and sex. This could be explained by the fact that sex differences are also known to exist in circadian rhythms ^71,72^ and in circadian genes rhythmicity between males and females in brains of humans^73^ and of rodents ^74^.

Our results are also consistent with prior studies which examined intersectional consequences between sex and circadian genes such as *Clock* or *Npas2* ^*44,75*^. Interestingly, *Npas2* deletion had higher impact on cocaine reward and self-administration behaviors in female mice ^44^. More profound consequences on females than males could be explained by levels of circulating hormones, as sex differences in cocaine self-administration were abolished in ovariectomized females ^44^. This was consistent with the fact that circulating estrogens were shown to be essential in orchestrating rhythms of circadian genes in the SCN ^76^. In addition, estrogen signaling has been shown to influence opioid tolerance and dependence behaviors ^77^. Thus, female circulating hormones could also be involved in the sexually dimorphic consequences of NPAS2 deficiency on opioid tolerance and dependence. Further studies examining the mechanisms of interaction between NPAS2, opioids sexual hormones are now warranted.

Overall, our study and prior studies examining the involvement of circadian genes deletion on opioid analgesia, tolerance, and dependence, show a differential involvement of *Npas2 and Per* genes ^29,30^. However, *Npas2* and *Per* genes bi-directionally influence each other’s expression levels, leading to variations in expression that follow circadian rhythmicity ^33^. Importantly, rhythmic expressions of circadian genes follow different circadian phases ^78,79^. Therefore, it is possible that rhythmic expression could have an impact on the involvement of each gene. In our study, we examined the role of NPAS2 deficiency at a similar time of day (ZT2) as the mPer1-KO and mPer2-KO studies (ZT3-5) ^29,30^. Ongoing studies are evaluating the impact of time of day and circadian genes on opioid-related behaviors.

It is important to note as well that while prior studies used morphine to evaluate the impact of *mPer1* and *mPer2* deletion, we used fentanyl to evaluate impact of NPAS2 deficiency. Although both opioids mediate analgesia via the mu-opioid receptor ^80,81^, they can activate different MOR downstream signaling pathways ^82-84^. We could therefore speculate that fentanyl and morphine may recruit different circadian genes and downstream signaling pathways involving the mu-opioid receptor.

In conclusion, our study provides evidence for a differential role of NPAS2 signaling in fentanyl mediated behaviors, with high impact on physical dependence and marginal effect on tolerance. Importantly, NPAS2 deficiency modulated these behaviors in a sexually dimorphic manner, with female mice more profoundly affected than males. Identification of NPAS2-controlled genes and signaling pathways that modulate opioid behaviors and that may interact with substrates of opioid tolerance and dependence, could provide better insight on understanding the impact of clock genes and circadian rhythmicity on the chronic use of prescription opioids in patients.

## REFERENCES

1. Hayhurst CJ, Durieux ME. Differential Opioid Tolerance and Opioid-induced Hyperalgesia: A Clinical Reality. Anesthesiology. Feb 2016;124(2):483–8. doi:10.1097/ALN.0000000000000963

2. Henry SG, Wilsey BL, Melnikow J, Iosif AM. Dose escalation during the first year of long-term opioid therapy for chronic pain. Pain Med. Apr 2015;16(4):733–44. doi:10.1111/pme.12634

3. Armenian P, Vo KT, Barr-Walker J, Lynch KL. Fentanyl, fentanyl analogs and novel synthetic opioids: A comprehensive review. Neuropharmacology. May 15 2018;134(Pt A):121–132. doi:10.1016/j.neuropharm.2017.10.016

4. Cao M, Javaheri S. Effects of Chronic Opioid Use on Sleep and Wake. Sleep Med Clin. Jun 2018;13(2):271–281. doi:10.1016/j.jsmc.2018.02.002

5. Garcia AN, Salloum IM. Polysomnographic sleep disturbances in nicotine, caffeine, alcohol, cocaine, opioid, and cannabis use: A focused review. Am J Addict. Oct 2015;24(7):590–8. doi:10.1111/ajad.12291

6. Mahfoud Y, Talih F, Streem D, Budur K. Sleep disorders in substance abusers: how common are they? Psychiatry (Edgmont). Sep 2009;6(9):38–42.

7. Sharkey KM, Kurth ME, Anderson BJ, Corso RP, Millman RP, Stein MD. Assessing sleep in opioid dependence: a comparison of subjective ratings, sleep diaries, and home polysomnography in methadone maintenance patients. Drug Alcohol Depend. Jan 15 2011;113(2-3):245–8. doi:10.1016/j.drugalcdep.2010.08.007

8. Bumgarner JR, Walker WH, 2nd, Nelson RJ. Circadian rhythms and pain. Neurosci Biobehav Rev. Oct 2021;129:296–306. doi:10.1016/j.neubiorev.2021.08.004

9. Eacret D, Veasey SC, Blendy JA. Bidirectional Relationship between Opioids and Disrupted Sleep: Putative Mechanisms. Mol Pharmacol. Oct 2020;98(4):445–453. doi:10.1124/mol.119.119107

10. Minett MS, Eijkelkamp N, Wood JN. Significant determinants of mouse pain behaviour. PLoS One. 2014;9(8):e104458. doi:10.1371/journal.pone.0104458

11. Warfield AE, Prather JF, Todd WD. Systems and Circuits Linking Chronic Pain and Circadian Rhythms. Front Neurosci. 2021;15:705173. doi:10.3389/fnins.2021.705173

12. Zhang P, Moye LS, Southey BR, et al. Opioid-Induced Hyperalgesia Is Associated with Dysregulation of Circadian Rhythm and Adaptive Immune Pathways in the Mouse Trigeminal Ganglia and Nucleus Accumbens. Mol Neurobiol. Dec 2019;56(12):7929–7949. doi:10.1007/s12035-019-01650-5

13. Hartwell EE, Pfeifer JG, McCauley JL, Moran-Santa Maria M, Back SE. Sleep disturbances and pain among individuals with prescription opioid dependence. Addict Behav. Oct 2014;39(10):1537–42. doi:10.1016/j.addbeh.2014.05.025

14. Huhn AS, Finan PH. Sleep disturbance as a therapeutic target to improve opioid use disorder treatment. Exp Clin Psychopharmacol. Jun 10 2021;doi:10.1037/pha0000477

15. Fathi HR, Yoonessi A, Khatibi A, Rezaeitalab F, Rezaei-Ardani A. Crosstalk between Sleep Disturbance and Opioid Use Disorder: A Narrative Review. Addict Health. Apr 2020;12(2):140–158. doi:10.22122/ahj.v12i2.249

16. Tripathi R, Rao R, Dhawan A, Jain R, Sinha S. Opioids and sleep - a review of literature. Sleep Med. Mar 2020;67:269–275. doi:10.1016/j.sleep.2019.06.012

17. Takahashi JS. Transcriptional architecture of the mammalian circadian clock. Nat Rev Genet. Mar 2017;18(3):164–179. doi:10.1038/nrg.2016.150

18. Hastings M. The brain, circadian rhythms, and clock genes. BMJ. Dec 19-26 1998;317(7174):1704–7. doi:10.1136/bmj.317.7174.1704

19. Rijo-Ferreira F, Takahashi JS. Genomics of circadian rhythms in health and disease. Genome Med. Dec 17 2019;11(1):82. doi:10.1186/s13073-019-0704-0

20. Naber D, Cohen RM, Pickar D, et al. Episodic secretion of opioid activity in human plasma and monkey CSF: evidence for a diurnal rhythm. Life Sci. Feb 23 1981;28(8):931–5. doi:10.1016/0024-3205(81)90056-4

21. Naber D, Wirz-Justice A, Kafka MS. Circadian rhythm in rat brain opiate receptor. Neurosci Lett. Jan 1 1981;21(1):45–50. doi:10.1016/0304-3940(81)90055-0

22. Petraglia F, Facchinetti F, Parrini D, Micieli G, De Luca S, Genazzani AR. Simultaneous circadian variations of plasma ACTH, beta-lipotropin, beta-endorphin and cortisol. Horm Res. 1983;17(3):147–52. doi:10.1159/000179690

23. Facchinetti F, D’Attoma G, Petraglia F, Pini LA, Sternieri E, Genazzani AR. Circadian variations of proopiocortin-related peptides in children with migraine. Cephalalgia. Aug 1983;3 Suppl 1:94–7. doi:10.1177/03331024830030S113

24. Vansteensel MJ, Magnone MC, van Oosterhout F, et al. The opioid fentanyl affects light input, electrical activity and Per gene expression in the hamster suprachiasmatic nuclei. Eur J Neurosci. Jun 2005;21(11):2958–66. doi:10.1111/j.1460-9568.2005.04131.x

25. Pacesova D, Volfova B, Cervena K, Hejnova L, Novotny J, Bendova Z. Acute morphine affects the rat circadian clock via rhythms of phosphorylated ERK1/2 and GSK3beta kinases and Per1 expression in the rat suprachiasmatic nucleus. Br J Pharmacol. Jul 2015;172(14):3638–49. doi:10.1111/bph.13152

26. Pacesova D, Novotny J, Bendova Z. The effect of chronic morphine or methadone exposure and withdrawal on clock gene expression in the rat suprachiasmatic nucleus and AA-NAT activity in the pineal gland. Physiol Res. Jul 18 2016;65(3):517–25. doi:10.33549/physiolres.933183

27. Cutler DJ, Mundey MK, Mason R. Electrophysiological effects of opioid receptor activation on Syrian hamster suprachiasmatic nucleus neurones in vitro. Brain Res Bull. Sep 15 1999;50(2):119–25. doi:10.1016/s0361-9230(99)00069-6

28. Li SX, Liu LJ, Jiang WG, Lu L. Morphine withdrawal produces circadian rhythm alterations of clock genes in mesolimbic brain areas and peripheral blood mononuclear cells in rats. J Neurochem. Jun 2009;109(6):1668–79. doi:10.1111/j.1471-4159.2009.06086.x

29. Perreau-Lenz S, Hoelters LS, Leixner S, et al. mPer1 promotes morphine-induced locomotor sensitization and conditioned place preference via histone deacetylase activity. Psychopharmacology (Berl). Jun 2017;234(11):1713–1724. doi:10.1007/s00213-017-4574-0

30. Perreau-Lenz S, Sanchis-Segura C, Leonardi-Essmann F, Schneider M, Spanagel R. Development of morphine-induced tolerance and withdrawal: involvement of the clock gene mPer2. Eur Neuropsychopharmacol. Jul 2010;20(7):509–17. doi:10.1016/j.euroneuro.2010.03.006

31. DeBruyne JP, Weaver DR, Reppert SM. CLOCK and NPAS2 have overlapping roles in the suprachiasmatic circadian clock. Nat Neurosci. May 2007;10(5):543–5. doi:10.1038/nn1884

32. Garcia JA, Zhang D, Estill SJ, et al. Impaired cued and contextual memory in NPAS2-deficient mice. Science. Jun 23 2000;288(5474):2226–30. doi:10.1126/science.288.5474.2226

33. Reick M, Garcia JA, Dudley C, McKnight SL. NPAS2: an analog of clock operative in the mammalian forebrain. Science. Jul 20 2001;293(5529):506–9. doi:10.1126/science.1060699

34. Puig S, Barker KE, Szott SR, Kann PT, Morris JS, Gutstein HB. Spinal opioid tolerance depends upon platelet-derived growth factor receptor-beta signaling, not mu-opioid receptor internalization. Mol Pharmacol. Jul 28 2020;doi:10.1124/mol.120.119552

35. Burma NE, Leduc-Pessah H, Trang T. Genetic deletion of microglial Panx1 attenuates morphine withdrawal, but not analgesic tolerance or hyperalgesia in mice. Channels (Austin).Jul 26 2017:1–8. doi:10.1080/19336950.2017.1359361

36. Stinus L, Le Moal M, Koob GF. Nucleus accumbens and amygdala are possible substrates for the aversive stimulus effects of opiate withdrawal. Neuroscience. 1990;37(3):767–73. doi:10.1016/0306-4522(90)90106-e

37. Puig S, Gutstein HB. Opioids: keeping the good, eliminating the bad. Nat Med. Mar 2017;23(3):272–273. doi:10.1038/nm.4277

38. Corder G, Tawfik VL, Wang D, et al. Loss of mu opioid receptor signaling in nociceptors, but not microglia, abrogates morphine tolerance without disrupting analgesia. Nat Med. Jan 16 2017;doi:10.1038/nm.4262

39. Zhou YD, Barnard M, Tian H, et al. Molecular characterization of two mammalian bHLH-PAS domain proteins selectively expressed in the central nervous system. Proc Natl Acad Sci U S A. Jan 21 1997;94(2):713–8. doi:10.1073/pnas.94.2.713

40. Zhang J, Li H, Teng H, et al. Regulation of peripheral clock to oscillation of substance P contributes to circadian inflammatory pain. Anesthesiology. Jul 2012;117(1):149–60. doi:10.1097/ALN.0b013e31825b4fc1

41. Ozburn AR, Falcon E, Twaddle A, et al. Direct regulation of diurnal Drd3 expression and cocaine reward by NPAS2. Biol Psychiatry. Mar 1 2015;77(5):425–433. doi:10.1016/j.biopsych.2014.07.030

42. Collett BJ. Opioid tolerance: the clinical perspective. Br J Anaesth. Jul 1998;81(1):58–68. doi:10.1093/bja/81.1.58

43. Gutstein HB AH. Opioid analgesics. In: Hardman JG LL, ed. Goodman and Gilman’s The Pharmacological Basis of Therapeutics. 10th ed. New York: McGraw-Hill ed. 2006:569.

44. DePoy LM, Becker-Krail DD, Zong W, et al. Circadian-Dependent and Sex-Dependent Increases in Intravenous Cocaine Self-Administration in Npas2 Mutant Mice. J Neurosci. Feb 3 2021;41(5):1046–1058. doi:10.1523/JNEUROSCI.1830-20.2020

45. Dudley CA, Erbel-Sieler C, Estill SJ, et al. Altered patterns of sleep and behavioral adaptability in NPAS2-deficient mice. Science. Jul 18 2003;301(5631):379–83. doi:10.1126/science.1082795

46. Gaudet AD, Fonken LK, Ayala MT, et al. Spinal Cord Injury in Rats Disrupts the Circadian System. eNeuro. Nov-Dec 2018;5(6)doi:10.1523/ENEURO.0328-18.2018

47. Schmidt BL, Tambeli CH, Barletta J, et al. Altered nucleus accumbens circuitry mediates pain-induced antinociception in morphine-tolerant rats. J Neurosci. Aug 1 2002;22(15):6773–80. doi:20026639

48. Johnson DW, Glick SD. Dopamine release and metabolism in nucleus accumbens and striatum of morphine-tolerant and nontolerant rats. Pharmacol Biochem Behav. Oct 1993;46(2):341–7. doi:10.1016/0091-3057(93)90362-w

49. Koob GF, Maldonado R, Stinus L. Neural substrates of opiate withdrawal. Trends Neurosci. May 1992;15(5):186–91. doi:10.1016/0166-2236(92)90171-4

50. Koob GF, Wall TL, Bloom FE. Nucleus accumbens as a substrate for the aversive stimulus effects of opiate withdrawal. Psychopharmacology (Berl). 1989;98(4):530–4. doi:10.1007/BF00441954

51. Parekh PK, Logan RW, Ketchesin KD, et al. Cell-Type-Specific Regulation of Nucleus Accumbens Synaptic Plasticity and Cocaine Reward Sensitivity by the Circadian Protein, NPAS2. J Neurosci. Jun 12 2019;39(24):4657–4667. doi:10.1523/JNEUROSCI.2233-18.2019

52. Georges F, Stinus L, Bloch B, Le Moine C. Chronic morphine exposure and spontaneous withdrawal are associated with modifications of dopamine receptor and neuropeptide gene expression in the rat striatum. Eur J Neurosci. Feb 1999;11(2):481–90. doi:10.1046/j.1460-9568.1999.00462.x

53. Pothos E, Rada P, Mark GP, Hoebel BG. Dopamine microdialysis in the nucleus accumbens during acute and chronic morphine, naloxone-precipitated withdrawal and clonidine treatment. Brain Res. Dec 6 1991;566(1-2):348–50. doi:10.1016/0006-8993(91)91724-f

54. Spanagel R, Almeida OF, Bartl C, Shippenberg TS. Endogenous kappa-opioid systems in opiate withdrawal: role in aversion and accompanying changes in mesolimbic dopamine release. Psychopharmacology (Berl). Jun 1994;115(1-2):121–7. doi:10.1007/BF02244761

55. Candido J, Lutfy K, Billings B, et al. Effect of adrenal and sex hormones on opioid analgesia and opioid receptor regulation. Pharmacol Biochem Behav. Aug 1992;42(4):685–92.

56. Cicero TJ, Nock B, Meyer ER. Gender-related differences in the antinociceptive properties of morphine. J Pharmacol Exp Ther. Nov 1996;279(2):767–73.

57. Kest B, Wilson SG, Mogil JS. Sex differences in supraspinal morphine analgesia are dependent on genotype. J Pharmacol Exp Ther. Jun 1999;289(3):1370–5.

58. Ali BH, Sharif SI, Elkadi A. Sex differences and the effect of gonadectomy on morphine-induced antinociception and dependence in rats and mice. Clin Exp Pharmacol Physiol. May 1995;22(5):342–4. doi:10.1111/j.1440-1681.1995.tb02012.x

59. Bartok RE, Craft RM. Sex differences in opioid antinociception. J Pharmacol Exp Ther. Aug 1997;282(2):769–78.

60. Craft RM, Stratmann JA, Bartok RE, Walpole TI, King SJ. Sex differences in development of morphine tolerance and dependence in the rat. Psychopharmacology (Berl). Mar 1999;143(1):1–7. doi:10.1007/s002130050911

61. Kest B, Palmese C, Hopkins E. A comparison of morphine analgesic tolerance in male and female mice. Brain Res. Oct 6 2000;879(1-2):17–22. doi:10.1016/s0006-8993(00)02685-8

62. Liu A, Zhang H, Qin F, et al. Sex Associated Differential Expressions of the Alternatively Spliced Variants mRNA of OPRM1 in Brain Regions of C57BL/6 Mouse. Cell Physiol Biochem. 2018;50(4):1441–1459. doi:10.1159/000494644

63. Holtman JR, Jr., Sloan JW, Wala EP. Morphine tolerance in male and female rats. Pharmacol Biochem Behav. Mar 2004;77(3):517–23. doi:10.1016/j.pbb.2003.12.020

64. Hoseini SM, Hosseini SA. Effect of dietary L-tryptophan on osmotic stress tolerance in common carp, Cyprinus carpio, juveniles. Fish Physiol Biochem. Dec 2010;36(4):1061–7. doi:10.1007/s10695-010-9383-x

65. Barrett AC, Cook CD, Terner JM, Craft RM, Picker MJ. Importance of sex and relative efficacy at the mu opioid receptor in the development of tolerance and cross-tolerance to the antinociceptive effects of opioids. Psychopharmacology (Berl). Nov 2001;158(2):154–64. doi:10.1007/s002130100821

66. Cicero TJ, Nock B, Meyer ER. Gender-linked differences in the expression of physical dependence in the rat. Pharmacol Biochem Behav. Jun 2002;72(3):691–7. doi:10.1016/s0091-3057(02)00740-2

67. Cepeda MS, Carr DB. Women experience more pain and require more morphine than men to achieve a similar degree of analgesia. Anesth Analg. Nov 2003;97(5):1464–8. doi:10.1213/01.ane.0000080153.36643.83

68. Sarton E, Olofsen E, Romberg R, et al. Sex differences in morphine analgesia: an experimental study in healthy volunteers. Anesthesiology. Nov 2000;93(5):1245-54; discussion 6A. doi:10.1097/00000542-200011000-00018

69. Fillingim RB, Gear RW. Sex differences in opioid analgesia: clinical and experimental findings. Eur J Pain. Oct 2004;8(5):413–25. doi:10.1016/j.ejpain.2004.01.007

70. Nasser SA, Afify EA. Sex differences in pain and opioid mediated antinociception: Modulatory role of gonadal hormones. Life Sci. Nov 15 2019;237:116926. doi:10.1016/j.lfs.2019.116926

71. Bailey M, Silver R. Sex differences in circadian timing systems: implications for disease. Front Neuroendocrinol. Jan 2014;35(1):111–39. doi:10.1016/j.yfrne.2013.11.003

72. Krizo JA, Mintz EM. Sex differences in behavioral circadian rhythms in laboratory rodents. Front Endocrinol (Lausanne). 2014;5:234. doi:10.3389/fendo.2014.00234

73. Lim AS, Myers AJ, Yu L, et al. Sex difference in daily rhythms of clock gene expression in the aged human cerebral cortex. J Biol Rhythms. Apr 2013;28(2):117–29. doi:10.1177/0748730413478552

74. Chun LE, Woodruff ER, Morton S, Hinds LR, Spencer RL. Variations in Phase and Amplitude of Rhythmic Clock Gene Expression across Prefrontal Cortex, Hippocampus, Amygdala, and Hypothalamic Paraventricular and Suprachiasmatic Nuclei of Male and Female Rats. J Biol Rhythms. Oct 2015;30(5):417–36. doi:10.1177/0748730415598608

75. Easton A, Arbuzova J, Turek FW. The circadian Clock mutation increases exploratory activity and escape-seeking behavior. Genes Brain Behav. Feb 2003;2(1):11–9. doi:10.1034/j.1601-183x.2003.00002.x

76. Hatcher KM, Royston SE, Mahoney MM. Modulation of circadian rhythms through estrogen receptor signaling. Eur J Neurosci. Jan 2020;51(1):217–228. doi:10.1111/ejn.14184

77. Bodnar RJ, Kest B. Sex differences in opioid analgesia, hyperalgesia, tolerance and withdrawal: central mechanisms of action and roles of gonadal hormones. Horm Behav. Jun 2010;58(1):72–81. doi:10.1016/j.yhbeh.2009.09.012

78. Harbour VL, Weigl Y, Robinson B, Amir S. Phase differences in expression of circadian clock genes in the central nucleus of the amygdala, dentate gyrus, and suprachiasmatic nucleus in the rat. PLoS One. 2014;9(7):e103309. doi:10.1371/journal.pone.0103309

79. Zhang R, Lahens NF, Ballance HI, Hughes ME, Hogenesch JB. A circadian gene expression atlas in mammals: implications for biology and medicine. Proc Natl Acad Sci U S A. Nov 11 2014;111(45):16219–24. doi:10.1073/pnas.1408886111

80. Matthes HW, Maldonado R, Simonin F, et al. Loss of morphine-induced analgesia, reward effect and withdrawal symptoms in mice lacking the mu-opioid-receptor gene. Nature. Oct 31 1996;383(6603):819–23. doi:10.1038/383819a0

81. Romberg R, Sarton E, Teppema L, Matthes HW, Kieffer BL, Dahan A. Comparison of morphine-6-glucuronide and morphine on respiratory depressant and antinociceptive responses in wild type and mu-opioid receptor deficient mice. Br J Anaesth. Dec 2003;91(6):862–70. doi:10.1093/bja/aeg279

82. von Zastrow M, Svingos A, Haberstock-Debic H, Evans C. Regulated endocytosis of opioid receptors: cellular mechanisms and proposed roles in physiological adaptation to opiate drugs. Curr Opin Neurobiol. Jun 2003;13(3):348–53.

83. Keith DE, Anton B, Murray SR, et al. mu-Opioid receptor internalization: opiate drugs have differential effects on a conserved endocytic mechanism in vitro and in the mammalian brain. Mol Pharmacol. Mar 1998;53(3):377–84.

84. Raehal KM, Schmid CL, Groer CE, Bohn LM. Functional selectivity at the mu-opioid receptor: implications for understanding opioid analgesia and tolerance. Pharmacol Rev. Dec 2011;63(4):1001–19. doi:10.1124/pr.111.004598

